# Generation of mutation hotspots in ageing bacterial colonies

**DOI:** 10.1101/041525

**Authors:** Agnieszka Sekowska, Sofie Wendel, Morten H. H. Nørholm, Antoine Danchin

## Abstract

How do ageing bacterial colonies generate adaptive mutants? Over a period of two months, we isolated on ageing colonies outgrowing mutants able to use a new carbon source, and sequenced their genomes. This allowed us to uncover exquisite details on the molecular mechanism behind their adaptation: most mutations were located in just a few hotspots in the genome and over time, mutations increasingly originated from 8-oxo-guanosine, formed exclusively on the transcribed strand.

## Introduction

Bacteria constitute a precious biological model system for studying the molecular details of ageing and evolution. Bacterial cells defective in the MutY enzyme, responsible for removing adenine nucleotides pairing with the 8-oxo oxidized variant of guanosine (8-oxo-G), exhibit a dramatic increase in the number of adaptive mutants and Bridges proposed a model to explain this phenomenon that was later termed retromutagenesis ^1,2^. In this model, the process of transcription opens up the DNA double helix, enhancing the probability of mutation in the transcribed strand, but only mutations on the transcribed strand are transferred to mRNA and translate into mutants proteins that explore novel activities. Subsequently, the activity of the mutant protein may enable the cell to replicate and thereby fixes the initial adaptive mutation on both DNA strands. In agreement with the retromutagenesis model, *lacZ* amber mutations on the transcribed strand were recently isolated in approximately 10fold excess over mutations on the non-transcribed strand upon treatment with the mutagen nitrous acid ^3^. However, the prevalence of this molecular mechanism has not been studied in a non-mutator background and has not been validated in a whole genome context.

To gain new, in-depth knowledge of the mechanisms behind adaptive mutation we wanted to study the genetic changes in a background as undisturbed as possible. For this purpose, we designed an *E. coli* strain incapable of fermenting maltose, plated it on rich medium with maltose, and subsequently collected all mutants starting to outgrow on colonies over the course of two months (Figure 1).

**Figure 1.**
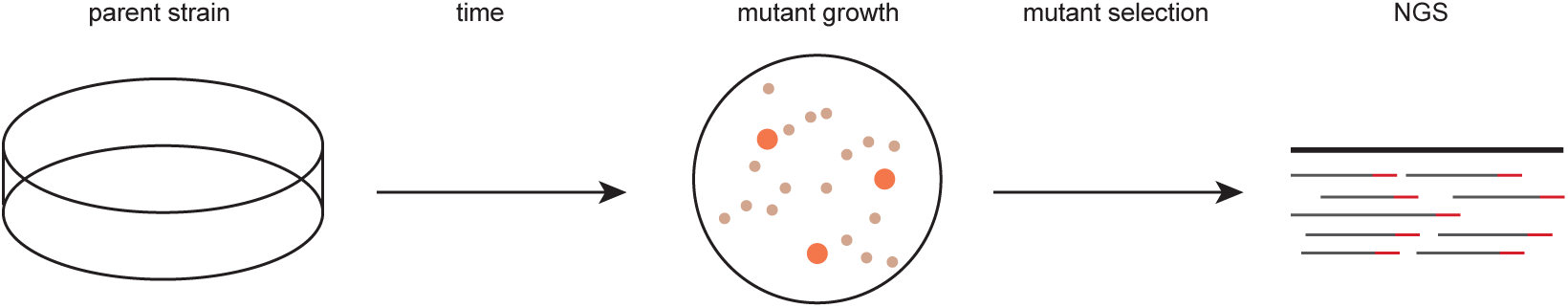
Schematic illustration of the experimental set-up. The parental strain is incapable of using the carbon source available in the plates, but after some days on plates, mutants adapted to their environment appear. These mutants are collected and subjected to next-generation genome sequencing (NGS).

## Results

### Aging colonies give rise to mutants in a non-mutator-based experimental system

Our starter strain is a derivative of the model bacterium *Escherichia coli* K12 MG1655 with its *cyaA* gene deleted, precluding synthesis of the signalling molecule cyclic AMP. As a consequence, a great many genes involved in carbon source catabolism cannot be expressed because they belong to operons requiring the cAMP Receptor Protein (CRP) complex to be activated ^4^. These cells can grow for some time on rich media, but, after having used the accessible sources present in the available medium (MacConkey medium), the cells cannot grow further but remain as small colonies caught in a quiescent state.

Around four days after plating, mutants capable of fermenting maltose started to appear as red papillae-like structures on the white colonies, and these mutants continued to appear over the next two months as some cells adapted to their environment. Mutant papillae outgrew on approximately one in 200 colonies, and progressively invaded the surface of the plate (Figure 2a) making it necessary to start with a large number of plates to be able to sample mutants at the late time points. All mutant strains were purified and displayed a variety of phenotypes on different carbon sources (Supplementary Table 1).

**Figure 2.**
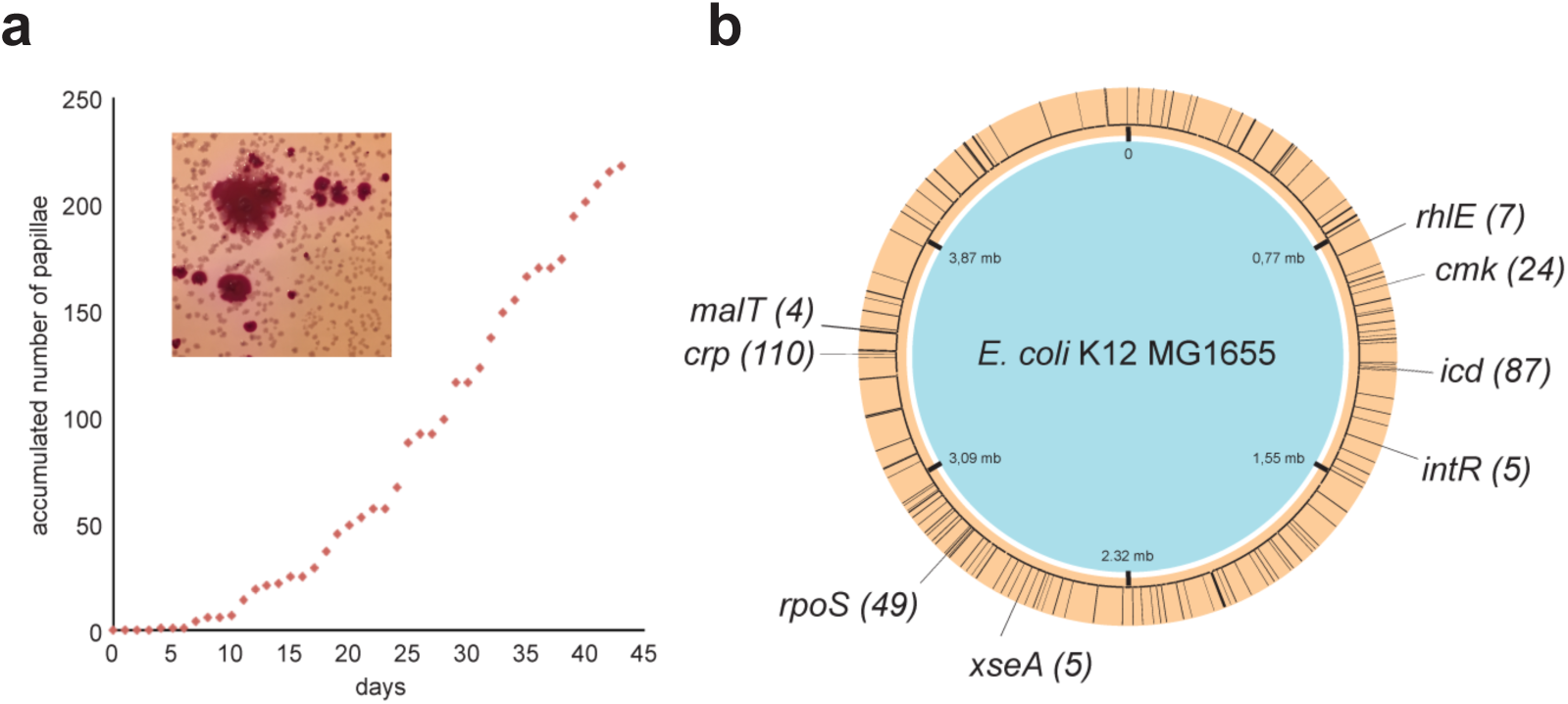
When subjected to prolonged incubation, *Escherichia coli* K12 MG1655 *cyaA*- cells adapt to their environment. (**a**) *E. coli cyaA*- cease to grow after forming small white colonies, when maltose is the major carbon source. After 4-5 days of incubation, mutants start occurring as red papillae on MacConkey agar (inserted image) and continue to form over approximately 6 weeks. (**b**) 96 mutant papillae were isolated, phenotypically characterized and genome sequenced. Loci with less than three mutations are uniformly distributed on the circular genome as illustrated with black lines. Hotspots of genes with more than two mutations are indicated with double-sized black lines and the total number of mutations in each gene is stated within brackets.

### Adaptive mutations are located in specific hotspot genes

We sequenced the genomes of 96 mutants, spanning the whole two-month period, and identified an average of four to five mutations per genome, the majority localized in a few hotspots only (Figure 2b and Supplementary Table 2). The hotspots are located in the following genes (Figure 2b): *cmk* (cytidylate kinase), *crp* (cyclic AMP receptor protein), *malT* (activator of the maltose regulon transcription), *rhlE* (ATP-dependent DNA helicase), *rpoS* (sigma factor sigma 38), *xseA* (large subunit of exonuclease VII), and two loci that are known to be unstable in laboratory strains; *e14-icd* (e14 prophage inserted in the NADP-dependent isocitrate dehydrogenase gene8) and *intR* (Rac prophage integrase9).

From the way the experiment was constructed, it could be expected that mutations in the *crp* gene would account for growth on maltose in a *ΔcyaA* background. However, the difficulty of obtaining cAMP-independent (*crp\**) mutants witnessed by Jon Beckwith’s laboratory suggested that under exponential growth such mutations were very rare (of the order of 1 in 10^9^ cells ^5^). In light of this, the ease with which we obtained such mutations in resting colonies was utterly unexpected.

It has been repeatedly observed that in *E. coli* laboratory strains, the *rpoS* gene is often inactivated ^6,7^. This is also what we observed at the *rpoS* hotspot (point mutations in general, but also frameshifts and a deletion of the region, see Supplementary Table 2). Inactivation of the RpoS protein may have triggered an increase in age-related mutagenesis, as this transcription factor is involved in oxidative stress response during stationary phase ^8^. However, before we understand the molecular mechanism, we can only speculate why the remaining hotspot genes are targeted.

### The majority of adaptive mutations take place on the transcribed strand

Theoretically, 12 different types of mutations are possible in DNA (A-C/G/T, C-A/G/T, G-A/C/T or T-A/C/G), but it is established that, due to respiration under stationary phase conditions, 8-oxo-G induced mutations are the most frequent to occur in the absence of an additional mutagenic process ^9,10^. In line with this, a dominant proportion of the mutations that were observed during the generation of the carbon-positive papillae is consistent with the involvement of 8-oxo-G (G-T and C-A transversions, 69% of the total number of missense mutations, Figure 3a). A large proportion of the remaining mutations (28% of the total number of missense mutations identified) are likely the result of cytosine deamination (G-C to A-T transitions, Figure 3a), another common mutational event in non-dividing cells ^9,11^.

**Figure 3.**
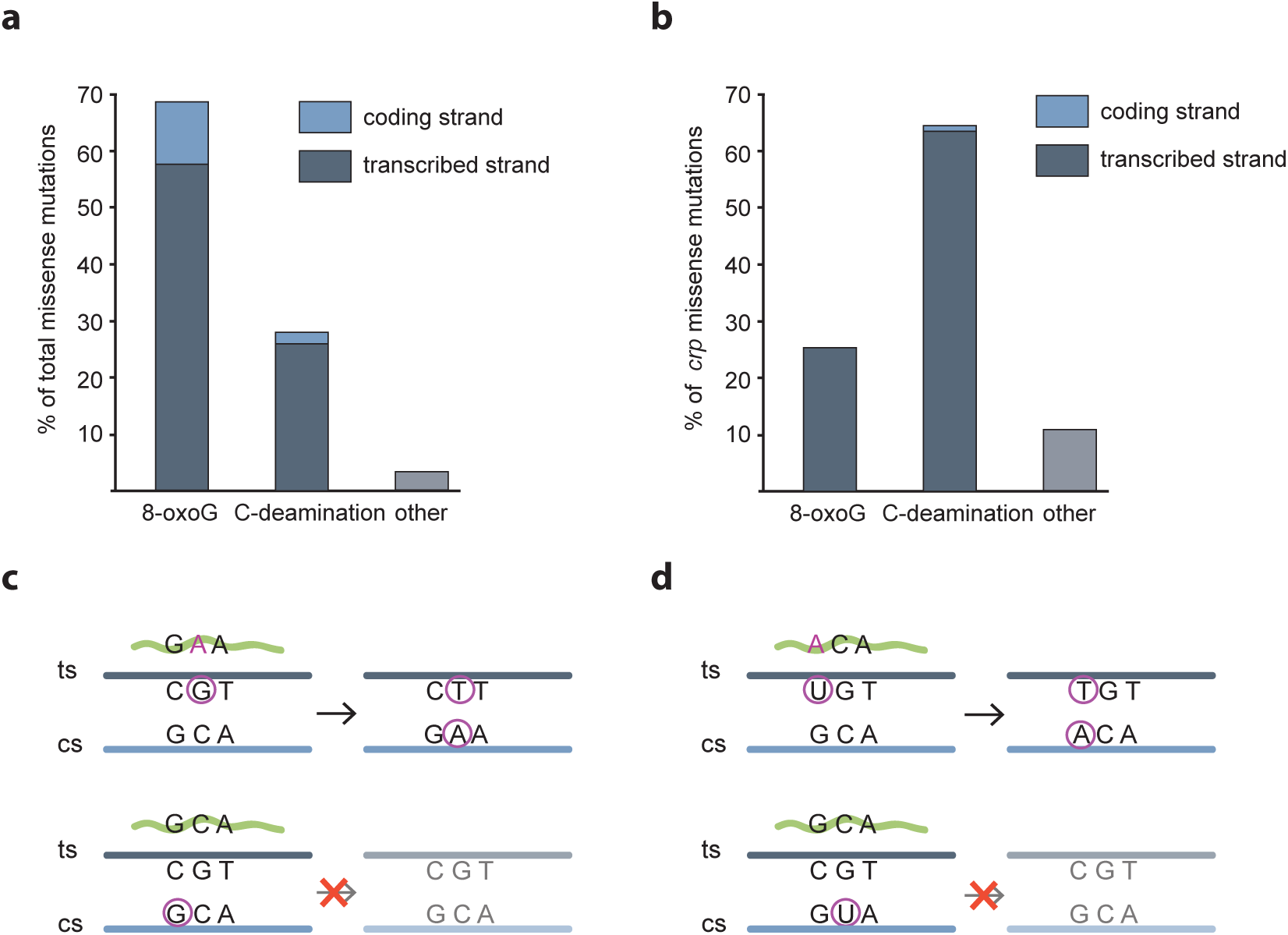
8-oxoguanine formation and cytosine deamination are highly dominating on the transcribed strand in isolated papillae. (**a**) Assuming that all identified G-T and C-A transversions are the result of 8-oxoguanine formation, and all G-C to A-T transitions are the result of cytosine deamination, we analysed the localisation of the corresponding G-and C-residues in the 96 sequenced mutant genomes and in (**b**) the 594 sequenced *crp* sequences. Only missense mutations were analysed. (**c**) Upper panel: Illustration of the mechanism of retromutagenesis after 8-oxoguanine (encircled G) formation on the transcribed strand (ts, dark grey line) that enables insertion of an adenine ribonucleotide in the mRNA (green waved line). The mutated mRNA enables growth and a round of replication that permanently fixes the mutation on the coding strand (cs, light blue line) in daughter cells. Lower panel: 8-oxoguanine formation on the coding strand does not transfer to daughter cells, (**d**) Illustration of retromutagenesis after cvtosine nucleotide deamination into uracil (encircled U). enabling base pairing with A. Upper and lower panels show result of cytosine deamination on transcribed and coding strand respectively, as detailed in (c).

Remarkably, the mutated base is highly dominantly present on the transcribed strand within gene coding sequences (84% for G transversions and 93% for C deaminations, Figure 3a). Even more extremely, in a total of 594 *crp* variants we sequenced from different papillae, 99% of the G transversion and C deamination events had taken place on the transcribed strand (Figure 3b). These observations are consistent with an increased mutation rate in transcribed regions (see Supplementary Table 2) and strongly support the retromutagenesis model (Figure 3 c-d) ^1,3,12^ – here in the absence of a mutagen. Importantly, the extreme strand bias considerably reduces the relevance of potential residual replication and growth in the aging colonies – a heavily debated, but hard to test, part of the controversy of adaptive mutations^13^.

In the retromutagenesis model, mutations generated on the transcribed strand are selected for. In contrast, a so-called hitchhiking mutation is per definition not selected. Are the other hotspots selected for or hitchhikers? It is interesting to compare the specific nucleotide changes in *crp* that occur alone (singles) with those found only in combination (paired) with a *crp\** mutation: 94% (399 out of 423) of single mutations were G transversion and C deamination events on the transcribed strand, whereas only 40% (27 out of 67) of the paired mutations were of this type. Furthermore, the 1% G transversions and C transitions found on the non-transcribed strand (eight events in total) were all in the paired positions. Thus, in contrast to the *crp\** mutations, it is not possible to decipher the mechanism leading to these extra mutations in *crp*.

C deamination events outside *crp* are evenly distributed (50% on the transcribed and the non-transcribed strand). This indicates that the other C deamination events are not caused by retromutagenesis. Apart from events in *crp*, G transversion mutations were identified in the *xseA*, *cmk*, *malT* and *rpoS* hotspots, and 20 out of 21 *cmk* and 5 out of 5 *xseA* G transversions had taken place on the transcribed strand. This may indicate a selective advantage of these mutants, but hitchhiking is still theoretically possible because the *xseA* and *cmk* genes are transcribed from the same (+) strand as *crp* on the genome and thus may be fixed in the same round of replication as the *crp\** mutants are. In contrast, *rpoS* mutations must have a selective advantage: 16 out of 17 *rpoS* G transversion missense mutations occur on the transcribed strand, but the *rpoS* gene is placed on the complement (-) strand and thus retromutagenesis must have taken place independently of *crp\** generation. Four out of four identified G transversion mutations in the *malT* hotspot were all placed on the non-transcribed strand and thus cannot have been selected by retromutagenesis.

### Formation of adaptive mutations shows age-dependence

Time is an important factor in the development of mutant papillae and the sampling of adaptive mutants over two months enabled us to observe interesting age-related trends. Firstly, a minimum of four days of incubation is required for adaptive mutations to occur in our experimental setup. Secondly, the frequency of papillae with *rpoS* mutations increases over time, suggesting an increasingly selective advantage with age (Figure 4a). Alternatively, the mechanism(s) leading to *rpoS* mutations are becoming more prominent over time. Thirdly, G transversion mutations in *crp* increases over time, paralleled by a decrease in C-deaminations (Figure 4b).

**Figure 4.**
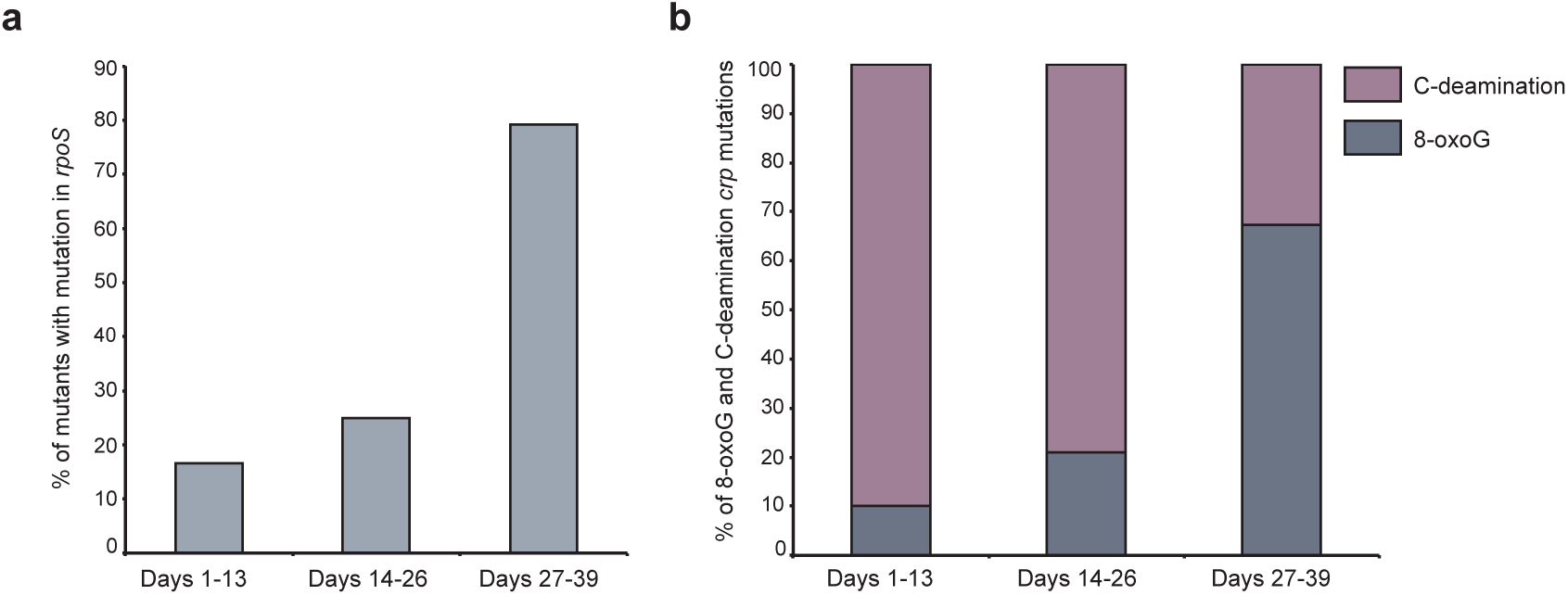
Time-dependent trends in mutational mechanisms. (**a**) The frequency of *rpoS* mutations in isolated mutants is increasing with time. The x-axis shows days after plating. (**b**) Isolated mutants show a distinct mutation pattern in the *crp* gene. Over time, there is a steady increase in 8-oxoG mutations whereas the C-deamination associated mutations decrease.

Our observations are not consistent with a general increase in mutagenesis, as the frequency of rifampicin resistance is of the order of four mutants per 10^8^ cells as generally observed ^14^. Furthermore, removal of RecA, a previously suggested key actor in adaptive mutations ^15^, displayed only a marginally negative effect on the total number of papillae formed (Supplementary Figure 1).

## Discussion

In the present study, we sequenced 96 adaptive mutants isolated during a two-month period, after having arisen from a minimally perturbed genetic background. To our knowledge, a similar experiment has not been performed at this time scale previously. Importantly, time was an essential factor not only for the development of adaptive mutation, but also seemingly for the mutational mechanism and we show that a highly dominating proportion of the adaptive mutations are 8-oxo-G modifications on the transcribed strand. This strongly supports the retromutagenesis model as a major mechanism behind adaptive mutations. Were this process generalised to multicellular organisms it would be a fertile contribution to the initiation of cancer that parallels ageing.

## Methods

Because the *fnr* gene is a homologue of *crp*, we used a starter strain previously identified as MG1655 at the ECGSC1 shown to harbour a ca. 40 kb deletion in the *fnr* region ^16^. The *cyaA* deletion was introduced with lambda Red-induced recombination ^17^. In pilot experiments, we had noticed that the presence of a pTrc99A plasmid (Pharmacia Biotech) carrying the *E. coli tig* gene (PCR amplified with the oligonucleotides CATGCCATGGTGAGGTAACAAGATGCAAGTTTC 3’ and 5‘ CGCGGATCCAATTACGCCTGCTGGTTCATC 3’ and cloned into the *Nco*I and *Bam*HI restriction sites) produced twice as many mutants that in absence of the *tig* plasmid, typically allowing us to recover around 200 mutants in each two months experiment, a figure quite convenient to get significant observations. Thus K12 Mg1655 (*cyaA*∷*cat* delta *fnr*, pTrc*tig*) is our starter strain. The *recA* deletion was from the Keio collections ^18^ that was used to prepare a P1 lysate by standard procedures.

For growth on plates the lactose-free MacConkey medium was used ^19^ (Difco MacConkey Agar Base) supplemented with 1% carbohydrate (maltose in most experiments) or glycerol and chloramphenicol, 5 mg/litre, ampicillin, 100 mg/litre, and 1 mM IPTG. For subsequent 3-times purification of papillae, chloramphenicol, ampicillin and IPTG were omitted from the plates. To obtain isolated colonies on the plates, early stationary-phase grown bacteria from the LB liquid culture (37°C) were diluted in sterile water containing 9 g/l sodium chloride, to the concentration of 2.5×10^4^ bacteria/ml and 100 μl of the bacterial suspension was spread onto MacConkey plates. The plates were subsequently placed in plastic boxes containing beakers with water to ensure constant humidity and placed for the duration of the experiment in an incubator at 37°C. To test for mutator phenotypes, rifampicin agar plates were prepared using the LB medium agar supplemented with 100 mg/litre rifampicin prepared in methanol. At all times the plates were wrapped in aluminium foil for protection from light.

The genomic libraries were generated using the TruSeq^^®^^Nano DNA LT Sample Preparation Kit (Illumina Inc.). 498 independent papillae were selected for amplification of the *crp* region by direct colony PCR. The PCR reaction was made using Red-Taq polymerase (Sigma-Aldrich) according to the manufacturer’s instructions (hybridisation at 55°C) with the following primers: forward primer 5’TTTCGGCAATCCAGAGACAGC3’ and reverse primer 5’AACATAGCACCAGCGTTTGTCG3’. The amplified *crp* regions were sequenced by Sanger method with two primers: forward 5’TTATCTGGCTCTGGAGAAAGC3’ and reverse primer 5’TCGAAGTGCATAGTTGATATCGG3’.

## Acknowledgements

This work was supported by The Novo Nordisk Foundation and a PhD grant from the People Programme (Marie Curie Actions) of the European Union’s Seventh Framework Programme [FP7-People-2012-ITN], under grant agreement No. 317058, “BACTORY”. We thank Ida Lauritsen and Emil Christian Fischer for technical assistance, and Anna Koza and Ida Bonde for their assistance with NGS.

## Author contributions

AD and MHHN designed the experiments, supervised the work and wrote the manuscript. AZ and SW carried out the experiments, contributed to the manuscript and prepared the figures.

## Competing financial interests statement

The authors have no competing financial interests.

